# Cationic inhalable particles for enhanced drug delivery to *M. tuberculosis* infected macrophages

**DOI:** 10.1101/2021.12.01.470866

**Authors:** Pallavi Raj Sharma, Ameya Atul Dravid, Yeswanth Chakravarthy Kalapala, Vishal K Gupta, Sharumathi Jeyasankar, Avijit Goswami, Rachit Agarwal

## Abstract

Inhalable microparticle-based drug delivery platforms are being investigated extensively for Tuberculosis (TB) treatment as they offer efficient deposition in lungs and improved pharmacokinetics of the encapsulated cargo. However, the effect of physical parameters of microcarriers on interaction with *Mycobacterium tuberculosis* (Mtb) infected mammalian cells is underexplored. In this study, we report that Mtb-infected macrophages are highly phagocytic and microparticle surface charge plays a major role in particle internalization by infected cells. Microparticles of different sizes (0.5 - 2 μm) were internalized in large numbers by Mtb-infected THP-1 macrophages and murine primary Bone Marrow Derived Macrophages *in vitro*. Drastic improvement in particle uptake was observed with cationic particles *in vitro* and in mice lungs. Rapid uptake of rifampicin-loaded cationic microparticles allowed high intracellular accumulation of the drug and lead to enhanced anti-bacterial function when compared to non-modified rifampicin-loaded microparticles. Cytocompatibility assay and histological analysis *in vivo* confirmed that the formulations were safe and did not elicit any adverse reaction. Additionally, pulmonary delivery of cationic particles in mice resulted in two-fold higher uptake in resident alveolar macrophages compared to non-modified particles. This study provides a framework for future design of drug carriers to improve delivery of anti-TB drugs inside Mtb-infected cells.

## 1. Introduction

Tuberculosis (TB) is a major health concern as it is the leading cause of death among all bacterial infectious diseases [1]. *Mycobacterium tuberculosis* (Mtb), the causative agent of TB, afflicted 10 million people in 2019, caused 1.2 million fatalities and an additional 208,000 deaths among HIV positive individuals [2]. TB therapy is demanding because the duration of treatment is long (6 – 24 months) with a high pill burden that leads to non-compliance among patients. Moreover, the emergence of antibiotic resistance is augmenting the issue with half a million patients reported with rifampicin (rif) resistance in 2019, of which 78% were multi-drug resistant [3].

Furthermore, effective drug delivery is challenging due to the unique pathogenesis of Mtb. Once inhaled, the bacteria deposit in the deep lung where they are phagocytosed by alveolar macrophages (AMs). Within these mammalian cells, Mtb survives as an intracellular pathogen and elicits host immune response that results in formation of a multi-cellular aggregate, called the granuloma, that contains the bacteria and infected cells [4]. The permeation of drugs inside the granuloma is variable. While drugs like rif are known to pervade through the diseased lung efficiently, others like moxifloxacin have poor permeation through the caseum of the granuloma [5]. Moxifloxacin, a fluoroquinolone antibiotic, has been shown to accumulate in immune cells *in vitro* and *in vivo* [6, 7]. The mechanisms utilized by fluoroquinolones to achieve high intracellular accumulation is not yet known, however, both passive and active mechanisms seem to be involved [8]. In granulomas, moxifloxacin concentration is correlated with the cellularity of the granuloma, with higher penetration into highly cellular granulomas [5]. In necrotic granulomas, moxifloxacin permeation is limited to the cellular rim, mostly localized within macrophages, and the drug does not diffuse through the necrotic caseous core [9]. Hence, to achieve therapeutic concentrations inside cells within the granuloma, high doses of anti-TB drugs are required for long duration which often cause systemic toxicity and side effects such as nausea, hepatoxicity, and nephrotoxicity [10]. There is a need to develop novel therapeutics and methods to improve efficiency of treatment to achieve high therapeutic concentrations at the site of infection, shorten treatment time and reduce adverse side-effects. To accomplish this, nano and micro particulate systems can be beneficial as drug delivery platforms [11]. Microparticles can be delivered through inhalation route to achieve targeted delivery to the lungs, which are the major site of infection in TB.

Sustained release formulation of drugs in polymeric microparticles (MPs) have significantly different pharmacokinetics and biodistribution compared to free drugs [12]. Anti-TB drugs are usually administered orally or intravenously, however, high local drug concentration of drug-loaded MPs can be achieved when drugs are delivered directly to the lung through inhalation [13, 14]. When delivered through inhalation route, drugs encapsulated in MPs exhibit longer serum half-life and bioavailability than an equivalent dose of the free drug because of superior deposition and retention in the lungs [15–17]. For pulmonary delivery, the optimal mass median aerodynamic diameter (MMAD) for microparticle deposition in the deep lung is < 5 μm [18]. Polymeric MPs in this size range (0.5 – 3 μm) are also phagocytosed efficiently by macrophages both *in vitro* and *in vivo* [19–21]. Encapsulating anti-TB drugs, such as rif, in such MP formulations has potential to improve pulmonary TB therapy. Once delivered to the lung, particulate delivery systems can be phagocytosed by cells and a higher intracellular concentration of drugs can be achieved.

Despite their advantages, MP-based platforms have not been translated to the clinic which may be attributed to the inability of the formulations to efficiently reach bacteria residing intracellularly [22]. Thus, it is imperative to understand how Mtb-infected cells interact with MPs and how such carriers can be engineered to improve delivery. Different materials and the physical and chemical properties of the formulation determine their interaction with different cell types, biodistribution, and clearance [23]. Surface charge is an important physical parameter that determines particle interaction with proteins and cells and its subsequent uptake by cells. Several studies have shown that cationic particles are safe and readily taken up by phagocytic cells both *in vitro* and *in vivo* [24–28]. However, the effect of varying surface charge of polymeric particles has not been studied with Mtb-infected macrophages. Given that Mtb infection has been known to modulate cytoskeletal proteins, cell membrane, phagocytic and endocytic machinery of the host, targeting strategies may differ between healthy and infected cells [29–32]. Through this study, we aim to design polymeric formulations that will be preferentially internalized by Mtb-infected cells and are optimal for *in vivo* delivery and uptake by immune cells through inhalation. We show that poly-l-lysine modified cationic microparticles exhibit higher uptake in immune cells both *in vitro* and *in vivo* compared to non-modified particles. Such modification also results in higher bactericidal efficacy and provide particle design principles for TB treatment.

## 2. Methods

### 2.1 Mice

Institutional Animal Ethics committee (CAF/Ethics/611/2018 and CAF/Ethics/792/2020) approval was obtained for conducting experiments on mice and IAEC guidelines were duly followed. C57BL/6 male mice were given standard food pellets and autoclaved water and were housed under constant temperature and humidity with automated 12 h light-dark cycles. Weight was regularly monitored and 8-10 weeks old male mice were used for experimentation. Ketamine (60 mg kg^-1^) and Xylazine (16 mg kg^-1^) were injected intraperitoneally to induce anaesthesia.

### 2.2 Bacterial culture

Mycobacterial strain H37Rv was handled in BSL-3 bacterial culture facility. Bacteria were cultured in Middlebrook 7H9 broth with 10% ADC (Bovine albumin fraction V 2.5 g, Dextrose 1 g, Catalase 0.0015 g in 50 ml distilled water), 0.5% Glycerol, and 0.05% Tween-80. For H37Rv-GFP, cultures were supplemented with Hygromycin (100 μg mL^-1^). The cultures were maintained at 37 °C with shaking at 180 rpm. For obtaining intracellular colony forming units (CFU), infected macrophages were permeabilized using 0.1% Triton-X followed by serial dilution with 1x PBS to relevant dilutions and plated on 7H11 plates supplemented with 10% OADC (Bovine albumin fraction V 2.5 g, Dextrose 1 g, Catalase 0.0015 g, Oleic acid 0.025 g in 50 ml distilled water). The plates were incubated at 37 °C and CFU were counted after 3-4 weeks of incubation as mycobacterial strains are slow growing.

### 2.3 THP-1 cell culture

THP-1 monocytic cells were maintained in RPMI media with 10% FBS at 37 °C and 5% CO2. Penicillin-Streptomycin was added to maintain mammalian cell culture but omitted for all bacterial infection experiments. All experiments were performed with cells between passage 3 and 18. Phorbol 12-myristate 13-acetate (PMA) stock (1 mg mL^-1^) was prepared in DMSO. For flow cytometry experiments, 2 x 10^5^ cells were seeded per well in a 24 well plate with 20 ng mL^-1^ PMA. After 18 h of incubation, cells were washed thrice with 1x PBS and fresh media was added. Differentiated macrophages were rested for 2 d after PMA treatment before further experimentation.

### 2.4 Microparticle synthesis

PLGA (Mw 10-15 kDa 50:50, Akina AP041) was used to synthesize microparticles using oil in water single emulsion method. Briefly, 100 mg of PLGA was dissolved in DCM for 10 min. For some experiments, Cy5 (25 μg mL^-1^) or rif (50 mg) were added to PLGA-DCM solution. This organic phase was added to 10 mL of 1% PVA and homogenized at 10,000 rpm (for 2 μm particles) and 12,000 rpm (for 1 μm particles) for 2 min. For 500 nm particles, probe sonication (Q125, Q-sonica) was used at 30% amplitude with 10 sec pulses for 2 min. The homogenized mixture was added to 100 mL of 1% PVA with constant stirring at 300 rpm for organic solvent evaporation. After 4-5 h, the particles were centrifuged at 10,000 g for 5 min and washed thrice with deionized water to remove residual PVA. The samples were snap-frozen and lyophilized (Taitec VD-250R) at −45 °C under vacuum. As PLGA will degrade in the presence of water, lyophilization is essential to remove residual water and allow long term storage of particles as powders. The particles were stored at −20 °C.

### 2.5 Macrophage infection with Mtb

Bacterial culture in log phase was pelleted at 5000 g for 10 min. The pellet was resuspended in RPMI media and passed through 26 G needle several times. This suspension was briefly sonicated and centrifuged at 300 g for 5 mins. The supernatant was collected in a fresh tube and OD at 600 nm was measured using spectrophotometer (Molecular Devices Spectramax M3). The bacteria were added to PMA differentiated THP-1 or BMDM cells at a multiplicity of infection of 10 and incubated for 4 h. After incubation, bacterial cells were washed away using 1x PBS followed by addition of media supplemented with Amikacin sulfate (200 μg mL^-1^) for 2 h, to kill the extracellular bacteria. Finally, cells were washed thrice with 1x PBS and incubated with fresh media at 37 °C and 5% CO2.

### 2.6 Rifampicin encapsulation

Rif (50 mg) was encapsulated in PLGA following the method described before to obtain rif loaded microparticles (rif-MPs) of 1 μm and 500 nm diameter. The amount of rif encapsulated was determined by dissolving 10 mg of particles in DMSO and measuring absorbance at 335 nm in a 96 well plate using an absorbance plate reader (Tecan Infinite Pro 2000). The absolute values were calculated using a standard curve of rif in 10 mg mL^-1^ PLGA dissolved in DMSO. The encapsulation efficiency was calculated by considering the ratio of the absolute value of rif in 100 mg PLGA and the theoretical maximum that can be encapsulated (50 mg).

### 2.7 Treatment of Mtb-infected macrophages with rif-MPs

Lyophilized particles (non-modified and PLL-conjugated blank and rif-loaded) were weighed and resuspended in sterile PBS (1 mg mL^-1^) and exposed to UV for 15 mins. Following brief sonication, particles were added to H37Rv-infected THP-1 macrophages at a final concentration of 50 μg mL^-1^. Free rif equivalent to encapsulated rif in particles was also added as a separate group. All wells were washed after 2 h exposure of particles or rif. After 72 h, all wells were washed thrice with 1x PBS and permeabilized with 0.1% Triton-X 100 for 10 mins. Intracellular bacteria obtained was serially diluted and plated on 7H11 agar plates and colonies were observed after 2-3 weeks.

### 2.8 Intratracheal delivery in mice

PLGA microparticles were resuspended in PBS (30 mg mL^-1^) and briefly sonicated. 50 μL of this suspension was loaded in a 3 mL syringe. A custom intratracheal tip with a 45° bent was attached to 20 G SS needle (Sigma Cat# Z101109) which was locked to the syringe.

Mice were anesthetized and mounted on a rodent stand at 45° angle. The tongue was pulled using sterile forceps and the chest was illuminated using a lamp to visualize the tracheal opening. An endotracheal tube was inserted into the trachea and the intratracheal tip was threaded through the tube into the trachea. The setup was kept stable and the syringe plunger was pushed to deliver PBS or particle suspension. The mouse was immediately dismounted, gently massaged, and allowed to recover on a heated pad.

### 2.9 Lung tissue preparation and immunostaining

After intratracheal delivery, mice were euthanized after 1.5 h. 30 mL of PBS was perfused through the heart to remove blood from the lung vessels. The lungs were dissected, chopped and treated with collagenase type-1 (5 mg mL^-1^) at 37 °C with constant rotation and intermittent mixing till the entire tissue was digested to release cells. The cell suspension was passed through 70 μm mesh followed by RBC lysis. The cells were resuspended in PBS-EDTA and stained with Fixable Aqua Live-Dead stain for 20 min at room temperature. Following one wash with FACS buffer (10% BSA in PBS-EDTA), the cells were fixed with 2% Paraformaldehyde (30 min, 4 °C) and stained with primary fluorophore conjugated antibodies for 30 min at 4 °C. Compensation beads were used for preparing single colour controls and relevant FMOs (Fluorophore minus one) were also stained. Finally, cells were resuspended in 300 μL FACS buffer and data was acquired using FACS DIVA and analysed using FlowJo (v9). Gating strategy to identify live immune cells, dendritic cells, alveolar and interstitial macrophages is explained in **Supplementary figure 13**.

### 2.10 *In vivo* imaging

Fluorescent PLGA particles were synthesised by encapsulating a near-IR dye (Cy7) to enable detection in a live animal using an *in vivo* imaging system (IVIS® Spectrum Perkin Elmer). Immediately after particle delivery, mice or explanted lungs were placed inside the instrument and X-ray images and fluorescence signal in Cy7 or Cy5 channel were acquired.

### 2.11 Statistical analysis

All statistical analyses were performed on GraphPad Prism 8.0.2. Values were represented as mean ± standard deviation (SD). Two tailed t-tests and one-way ANOVA were used where applicable and *p* < 0.05 was considered significant. One-way ANOVA was used to assess significance among two or more groups followed by Tukey’s test for post-hoc analysis which is recommended when all possible pairwise comparisons are required.

## 3. Results

### 3.1 PLGA microparticles synthesized with varying size and charge

Poly (lactic-co-glycolic) acid (PLGA) microparticles of different sizes were synthesized using oil in water single emulsion method, which is a simple and adaptable method to synthesize microparticles with tuneable sizes and allows encapsulation of hydrophobic drugs and dyes. Particles in the size range 500 nm - 3 μm are considered optimum for phagocytosis [19, 24, 33]. We synthesized particles with diameter 500 nm, 1 μm and 2 μm to understand the particle uptake behaviour of each sized microparticle by macrophages. Fluorescent dye, Cy5, was incorporated in the organic phase during synthesis to visualize these particles (**Figure 1A**). Size distribution was determined using DLS (**Figure 1B**) and showed a low polydispersity index of around 0.2 (**Table 1**). Particle surface morphology was smooth and spherical, as evident from SEM images (**Supplementary figure 1**). To test the cytocompatibility of the particle formulations, WST-8 assay was performed with THP-1 macrophages. As previously reported for PLGA particles, concentrations tested at 50 μg mL^-1^ and 500 μg mL^-1^ were found to be cytocompatible (**Supplementary figure 2**). A particle concentration of 50 μg mL^-1^ was used for all subsequent experiments.

**Figure 1.**
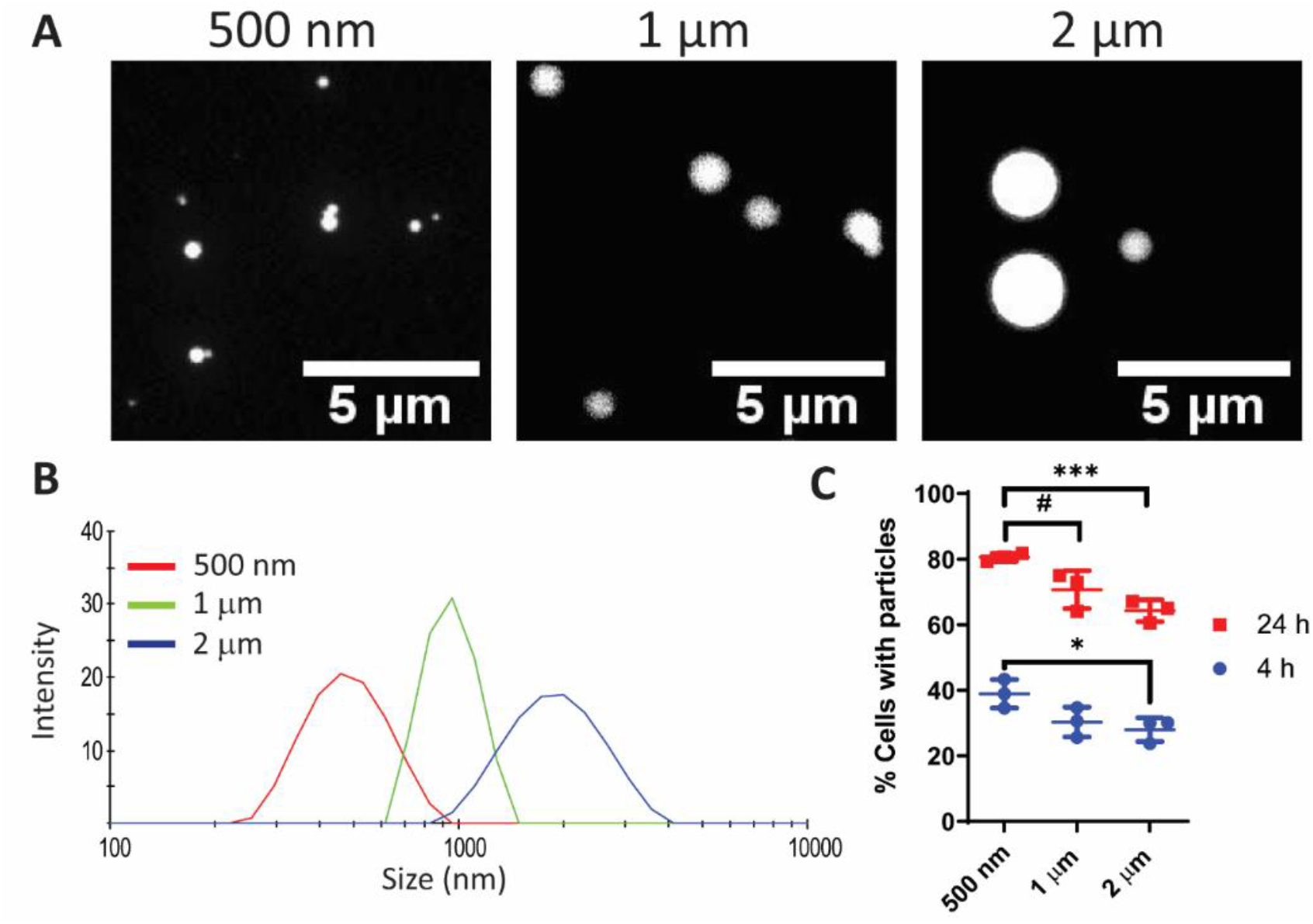
PLGA Microparticles of different sizes are readily phagocytosed by macrophages. (A) Representative fluorescence images of microparticles of sizes 500 nm,1 μm, and 2 μm. Images were acquired in the Cy5 channel under 100x magnification. Scale 5 μm. (B) Size distribution of microparticles was determined using Dynamic Light Scattering. (C) Percentage of differentiated THP-1 macrophages with particles over 4 and 24 h. One-way ANOVA followed by Tukey’s multiple comparison test was used to detect statistical differences for each time point (\**p* = 0.0423, #*p* = 0.0487, \*\*\**p* = 0.0053, n = 3). Data were represented as mean ± SD.

**Table 1:** Size and charge characterization of synthesized PLGA microparticles. PLL = Poly-l-lysine

| Target<br>Particle size | Average<br>Diameter (nm) | Polydispersity<br>index | Zeta potential<br>(mV) | Zeta potential<br>PLL conjugated (mV) |
| --- | --- | --- | --- | --- |
| 500 nm | 505 | 0.216 | -21.53 ± 5.14 | 21.73 ± 3.1 |
| 1 µm | 1198 | 0.231 | -23.43 ± 2.47 | 23.16 ± 3.34 |
| 2 µm | 2192 | 0.18 | -26.40 ± 3.73 | 24.93 ± 3.06 |

Next, to examine the effect of microparticle size, we studied uptake of these particles by macrophages *in vitro.* THP-1 macrophages were incubated with particle formulations and at specific time points, cells were washed thrice to remove adsorbed particles and analysed using flow cytometry (**Supplementary figure 3**). It was observed that 500 nm particle had slightly higher uptake as compared to 1 μm and 2 μm microparticles at both 4 h (38.9 ± 3.5% of cells, *p* = 0.0423) and 24 h (80.5 ± 0.98% of cells) post incubation (**Figure 1C)**. This could be due to higher particle to cell ratio in case of 500 nm particles (1 mg of 500 nm particles has a greater number of particles than 1 mg of 1 μm particles). Microparticles of size 1 μm and 2 μm were also phagocytosed by ~30% cells in 4 h and by ~65% cells by 24 h. Overall, we did not observe any major increase in the percentage of cells taking up particles of different sizes suggesting that all particles in these size ranges have similar uptake by macrophages.

### 3.2 Positive surface charge contributes to enhanced uptake by macrophages

Next, we explored the effect of particle surface charge on particle uptake by macrophages. The synthesized particles had a negative surface charge due to the presence of carboxylic acid end groups in the PLGA polymer (**Table 1**). Carboxylic acid end groups are deprotonated at neutral pH and results in negative charge on the particles. We covalently conjugated a US FDA approved cationic lysine homopolymer, poly-L-lysine (PLL), to particle surface using EDC-NHS chemistry. The zeta potential measurements confirmed the positive surface charge on the particles (**Table 1**).

When THP-1 macrophages were incubated with poly-L-Lysine (PLL) conjugated microparticles (PLL-MPs), the uptake of microparticles was enhanced two-fold compared to non-modified microparticles. For all sizes of microparticles, ~60% of macrophages had internalized positively charged particles in 4 h compared to <40% for non-modified particles (**Figure 2A**). Additionally, an enhanced Median Fluorescence Intensity (MFI) was observed with positively charged particles, indicating that along with the number of cells that had internalized particles, the number of particles being internalized per cell had also increased significantly (**Supplementary figure 4)**. By 24 h, both non-modified and positively charged particles were internalized comparably by the macrophages **(Supplementary figure 5)**. These results suggest that positively charged particles are rapidly taken up by macrophages possibly due to the increased coulombic attraction between cationic particles and anionic cell surface. The internalization of particles was further confirmed using confocal microscopy, where nearly all particles were observed inside the cells (**Figure 2C**).

**Figure 2.**
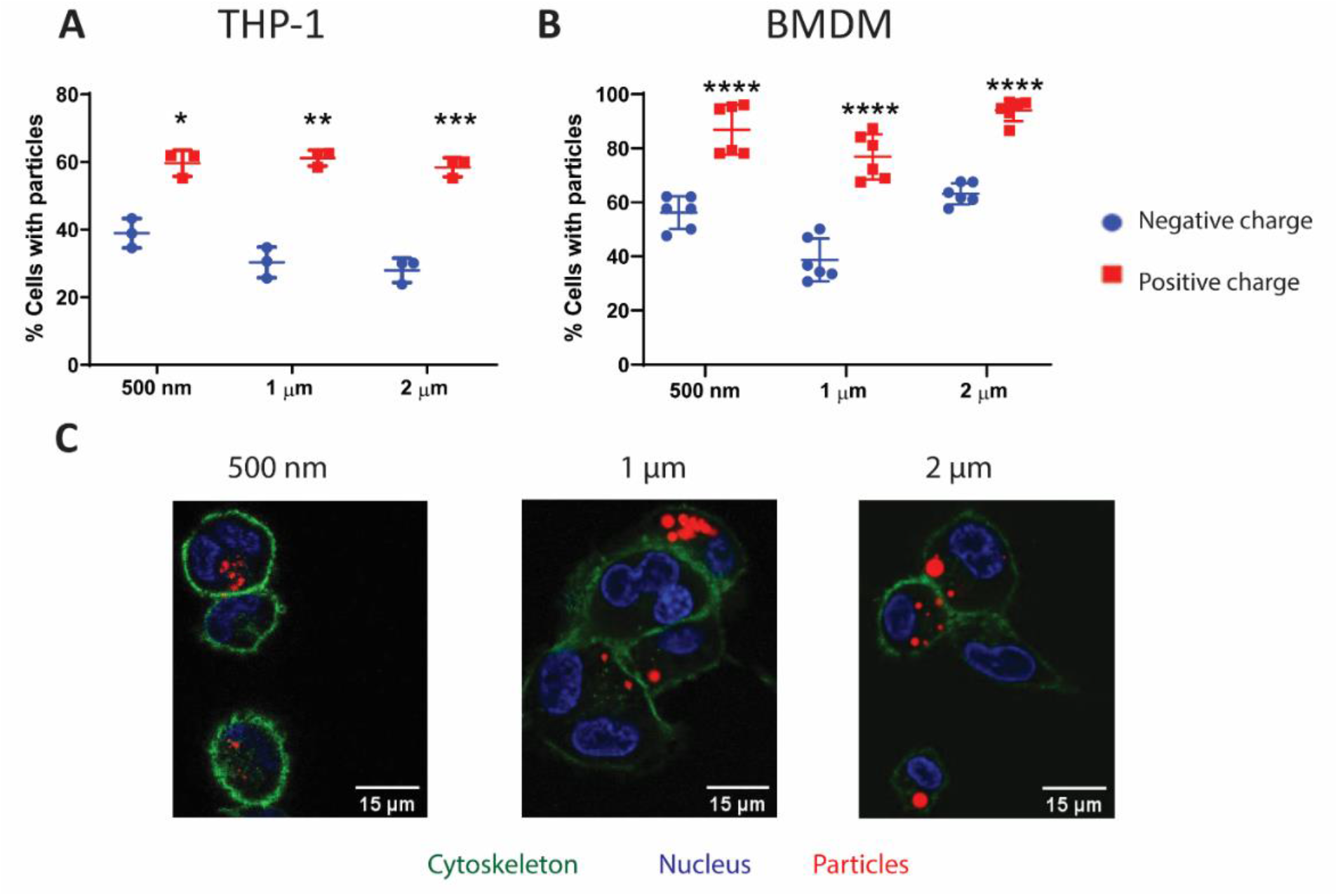
Cationic microparticles exhibit rapid uptake by macrophages. (A) Effect of particle surface charge on uptake by THP-1 macrophages after 4 h of incubation (n =3, \**p* = 0.0048, \*\**p* = 0.0002, \*\*\**p* = 0.001) (B) Effect of particle size and charge on particle uptake by primary murine BMDMs after 2 h incubation (n = 6, N = 2, \*\*\*\**p* < 0.0001) (C) Representative fluorescence microscopy images of THP-1 macrophages with positively charged microparticles of different sizes. Nucleus = Blue, Particles = Red, Cytoskeleton (Actin) = Green. For (A) and (B), statistical analysis was done using ordinary one-way ANOVA followed by Tukey’s multiple comparison test for each time point, two-tailed t-test was used to compare between non-modified and PLL-MPs. Data were represented as mean ± SD.

To confirm whether these effects were cell line specific or were also applicable on primary cells, we used mouse BMDMs and analysed CD11b and F4/80 surface marker expression to confirm purity (**Supplementary figure 6**). Upon incubation with all sizes of microparticles for 2 h, non-modified particles showed similar uptake as was the case with THP-1 **(Figure 2B)**. When incubated with PLL-MPs, enhanced particle-positive population was observed for each size (*p* < 0.0001) (**Figure 2B**) along with higher MFIs (**Supplementary figure 7**). Almost 90% of BMDMs phagocytosed the PLL-MPs within just 2 h of incubation indicating higher uptake potential of primary BMDMs.

### 3.3 Mtb-infected macrophages phagocytose cationic microparticles rapidly

While several studies have looked at the effect of particle uptake in healthy cells, the effect of Mtb infection on its uptake is still sparsely explored. To verify our observations in an *in vitro* infection model, we moved on to study how particle size and charge affects phagocytosis by Mtb-infected macrophages. In H37Rv-infected THP-1 macrophages (**Figure 3A**) and H37Rv-infected BMDMs, 500 nm and 1 μm particles were most efficiently phagocytosed (**Figure 3B**). We again observed a decrease in particle uptake with increasing particle size which can be attributed to a higher particle to cell ratio. Like non-infected macrophages, infected macrophages showed much higher uptake of PLL-MPs, with ~90% H37Rv-infected cells with particles within 4 h of incubation for each size. The highest uptake was observed by 500 nm and 1 μm positively charged particles in both THP-1 and BMDMs (**Figure 3A-B**). Corresponding enhancement in MFI was also observed for infected cells that had internalized particles (**Supplementary figure 8 and 9**). Another interesting observation across all infected groups was that bacteria-harbouring macrophages (GFP+ cells) had a higher Cy5+ population compared to cells that were exposed to bacteria but had not taken them up (uninfected macrophages) (**Supplementary figure 10**). This indicates that macrophages infected with virulent Mtb have high phagocytic ability. Similar results were demonstrated previously with *M. bovis* BCG infected rat AMs and *M. tuberculosis* infected THP-1 and *ex vivo* cultured primary murine lung macrophages [31, 32]. This suggests that the use of particles could also provide better targeting to infected cells. The higher uptake of PLL-MPs was confirmed using confocal microscopy with BMDMs infected with H37Rv after 2 h of incubation with particles (**Figure 3C**). Increased colocalization of bacteria and particles was also observed (indicated by white arrows in **Figure 3C**).

**Figure 3.**
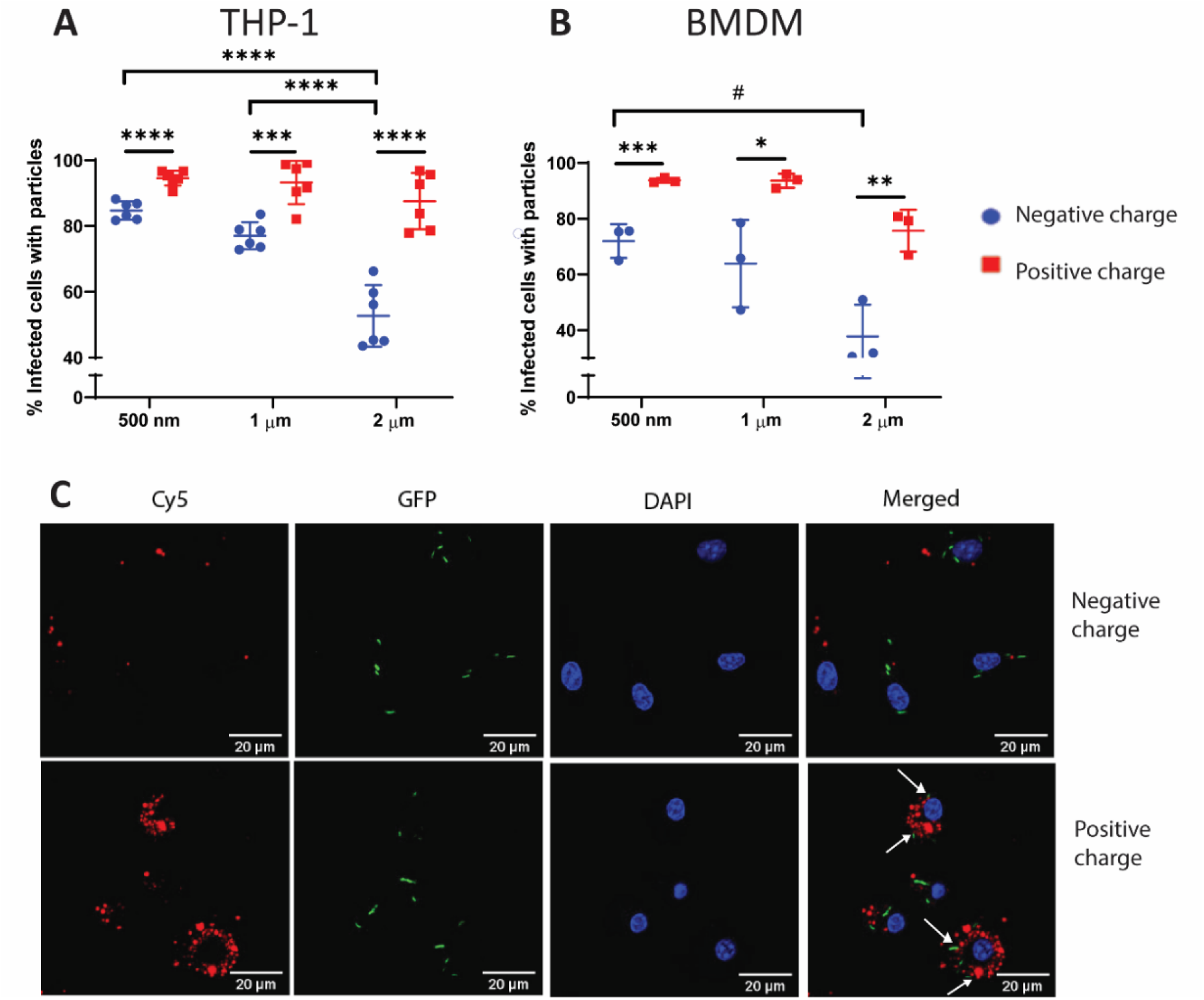
Cationic microparticles demonstrate higher uptake in Mtb-infected macrophages and colocalize with intracellular bacteria. Effect of surface charge on microparticle uptake in (A) H37Rv-infected THP-1 macrophages after 4 h incubation (\*\*\**p* = 0.0005, \*\*\*\**p* < 0.0001, n = 6, N = 2) (B) H37Rv-infected murine BMDMs after 2 h incubation (\*\*\**p* = 0.0035, \*\**p* = 0.0087, \**p* = 0.0316, #*p* = 0.0273, n = 3) (C) Representative confocal fluorescence images of 500 nm negatively and positively charged particle uptake in H37Rv-infected BMDMs. Nucleus = Blue, Bacteria = H37Rv-GFP green, Particles = Cy5 red. White arrows indicate instances of colocalization between particles and bacteria. Scale bar 20 μm. Statistical test conducted was two tailed t-test between the negative and positive charge groups for each size and one-way ANOVA followed by Tukey’s multiple comparison test for analysis among sizes. Data were represented as mean ± SD.

### 3.4 Enhanced bactericidal activity of cationic rif-MPs against intracellular Mtb

To test if this charge dependent effect has a functional relevance and can be applied to improve drug delivery in Mtb infected cells, we encapsulated rif in a 1 μm PLGA particle and were able to load 60 μg per mg particle. Rif encapsulation did not affect particle morphology, and particles remained smooth and spherical (**Figure 4A**). Drug release was assessed in PBS at 37 °C with continuous shaking over a few days and it was observed that ~40% of the drug is released within 3 d (**Figure 4B**). This is because we used a low molecular weight acid end capped PLGA with lactide to glycolide ratio 50:50 for microparticle synthesis which results in relatively faster degradation and drug release. Release was also performed at pH 5.5 to mimic conditions present in acidified phagosomes, where the MPs might get localized after phagocytosis. We found that lowering the pH accelerates the release of the encapsulated drug (**Figure 4B**). This could be because of the elevated rate of acid catalysed hydrolysis of the polymers at lower pH. Both formulations showed sustained release of rif for over 3 weeks (**Figure 4B**). The surface charge on these rif loaded MPs was modified by conjugating poly-L-lysine as previously described. The zeta potential was transformed from −24.73 ± 0.7 mV to +20.73 ± 0.6 mV. The drug encapsulation had a negligible difference before and after EDC-NHS conjugation of PLL.

**Figure 4.**
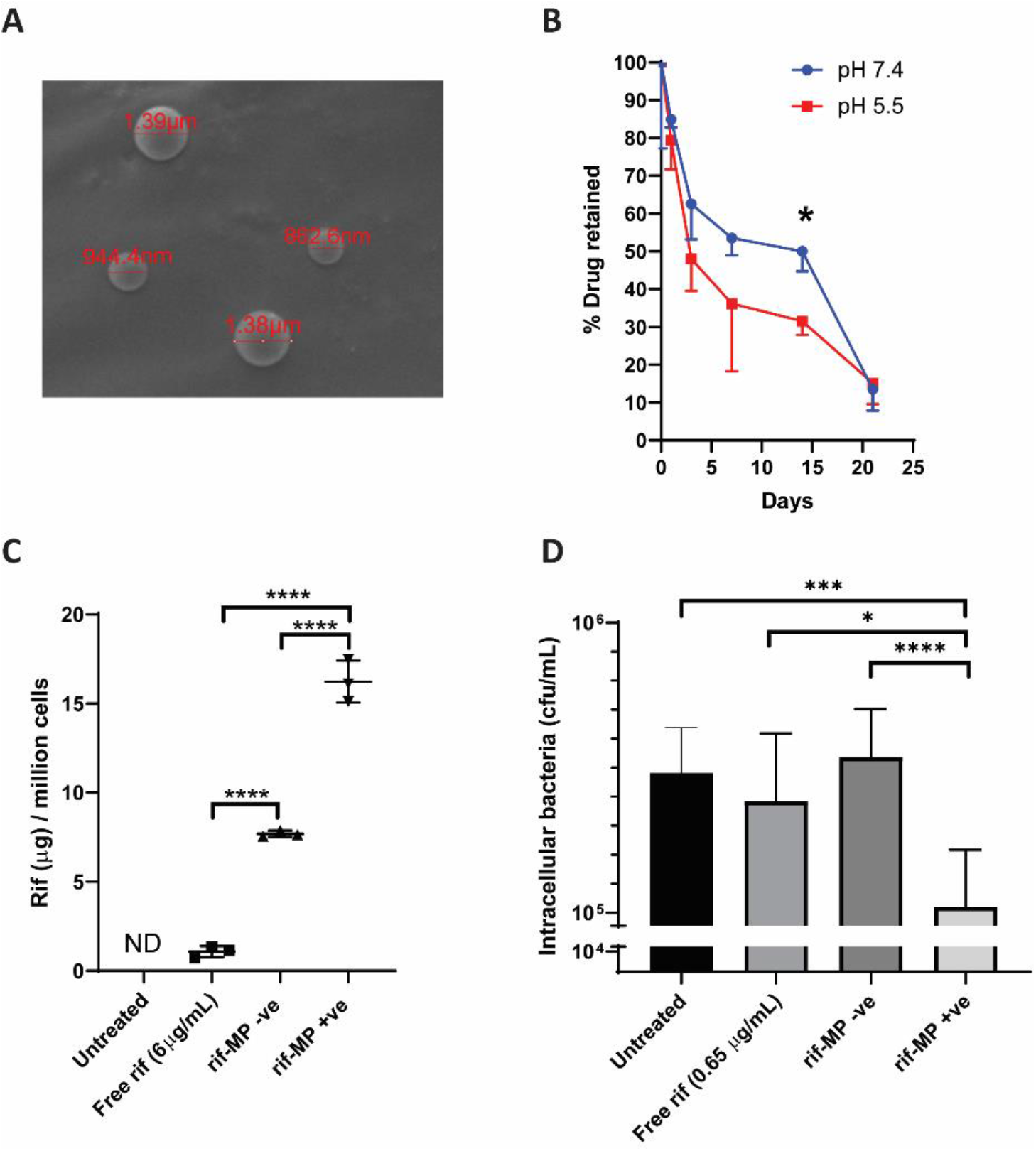
Cationic rif-microparticles delivered higher drug amounts in macrophages and were effective in the clearance of intracellular Mtb. (A) Representative SEM image of rif-loaded 1 μm PLGA particle with PLL coating (B) Percentage of drug retained in microparticles when incubated in 1x PBS (pH 7.4) and 0.1 M MES (pH 5.5) over 21 d (n =3, \**p* = 0.0076) (C) Intracellular rif quantification after 4 h treatment with free rif and rif-MPs (n = 3, \*\*\*\**p* <0.0001, ND = Not Detectable) (D) Intracellular bacteria from H37Rv infected THP-1 macrophages after 2 h treatment with free rif and encapsulated rif-MPs (n = 16, N = 2, \*\*\*\**p* <0.0001, \*\*\**p* = 0.0007, \**p* = 0.0314). Data were combined from two separate biological replicates. For (B), two-tailed t-test was applied on each time point to assess significance. For (C) and (D), ordinary one-way ANOVA followed by Tukey’s multiple comparisons test was used to determine significance. Data were represented as mean ± SD.

To determine if positively charged particles can deliver higher drug amounts in cells compared to negatively charged particles and free drug, THP-1 macrophages were incubated with MPs (100 μg mL^-1^). After 4 h of incubation, the cells were washed, lysed and intracellular rif content was quantified using HPLC. It was observed that rif-MPs encapsulating equivalent rif concentration allowed a log-fold higher amount of rif to accumulate within cells compared to free drug (**Figure 4C**). PLL-coated rif-MP further resulted in ~2-fold higher intracellular rif amount compared to non-modified rif-MP (*p* <0.0001) (**Figure 4C**). Thus, rif loaded microparticles create an intracellular reservoir of the drug and high concentrations can be achieved within infected cells. To test bactericidal efficacy inside mammalian cells, THP-1 macrophages were infected with H37Rv followed by 2 h incubation with free rif (0.65 μg mL^-1^) as well as both negative and positive surface charge rif-MPs with equivalent amount of rif. Only PLL-rif MP were able to reduce the intracellular bacterial counts after 72 h compared to untreated (*p* = 0.0007), free rif (*p* = 0.0314) and non-modified rif-MP (*p* <0.001), which indicates that prompt uptake of PLL-coated MPs leads to higher accumulation of the intracellular drug and higher bactericidal efficiency (**Figure 4D**). Blank microparticles of both surface charge had no effect on intracellular bacterial counts (**Supplementary figure 11**). Thus, PLL-MPs represent an improved drug delivery platform to target and treat Mtb-infected macrophages *in vitro*.

### 3.5 PLL-MPs show enhanced uptake in immune cells *in vivo*

Next, we studied if the rapid uptake of PLL-MPs would also be relevant *in vivo*. Non-modified and PLL-MPs were delivered to mice lungs via intratracheal route. Particles were suspended in PBS and 50 μL of this suspension (1.5 mg per mouse) was aerosolised into the mouse lung directly through the trachea. The particles were found to be localized to the lungs (**Supplementary figure 12A and 12B**) and were uniformly distributed (**Figure 5A).** Particles were also observed in the alveolar space **(Figure 5B**) suggesting deep lung delivery. Safety of the formulation was determined by histological analysis 1 and 7 d after particle administration, which revealed no undesired pathology in mice lungs administered with PBS, non-modified and PLL-MPs (**Figure 5E**).

**Figure 5.**
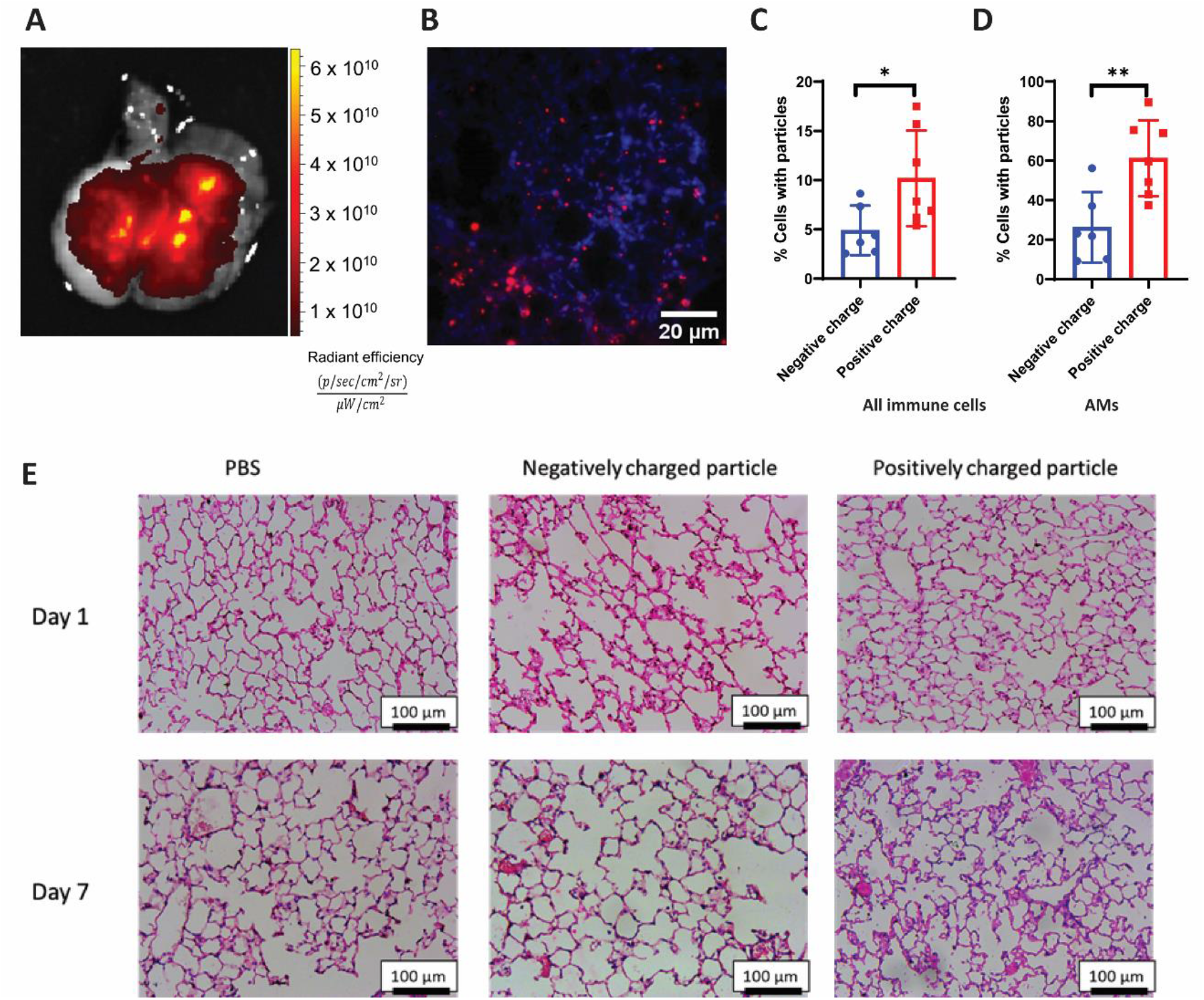
Cationic microparticles exhibit enhanced uptake by immune cells in vivo after pulmonary delivery and do not elicit inflammatory reactions. (A) Cy5-loaded MPs were delivered intratracheally and fluorescence intensity was measured immediately post-delivery from excised lung. (B) Microparticles deposition and distribution throughout the lung was assessed by fluorescence imaging of lung cryosections. Red = Cy5, Blue = Hoechst 33342, Scale bar = 10 μm. Percentages of cells with particles reported for (C) total immune cells (\**p* = 0.0356) and (D) alveolar macrophages (AMs) (\*\**p* = 0.0062) characterized using flow cytometry. Two-tailed t-test was used to determine significance between the two groups. Data were represented as mean ± SD. (E) Representative brightfield images of Hematoxylin and Eosin-stained lung sections after PBS and particle (negative and positive charge) exposure via intra-tracheal delivery. Mice were sacrificed after 1 and 7 d post-delivery. Scale bar = 100 μm.

Next, we proceeded with studying the effect of surface charge on particle uptake by immune cells *in vivo.* After 1.5 h of particle delivery, mice were euthanized and lungs were excised and digested with collagenase. Various immune cell populations (CD45+) such as neutrophils (CD11b+ Ly6G+), dendritic cells (CD24+ CD11c+), alveolar macrophages (CD24-CD11b-SiglecF+) and interstitial macrophages (CD24-CD11b+ SiglecF-) were identified using flow cytometry and particle uptake by each cell type was analysed (**Supplementary figure 13**). Particle delivery to immune cells and AMs was significantly enhanced with PLL-MPs. For all immune cells, non-modified particles were taken up by 4.89 ± 2.3% cells while PLL-MPs exhibited two-fold higher uptake with 10.18 ± 4.5% cells with particles (*p* = 0.0356) (**Figure 5C**). Similarly for alveolar macrophages, drastic improvement in delivery was observed with the percentage of cells with particles increasing from 26.25 ± 16.27% to 61.18 ± 17.78% with PLL-MPs (*p* = 0.0062) (**Figure 5D**). Particle uptake in other immune cell subsets also followed a similar trend with PLL-MPs (**Supplementary figure 14**). Thus, PLL-MPs represent an effective platform to enhance delivery to alveolar macrophages.

## 4. Discussion

In our study, mycobacteria infected macrophages were found to be highly phagocytic towards PLGA microparticles *in vitro,* which has been reported before in rat AMs [31]. Particle uptake was remarkably enhanced by conjugating with poly-L-lysine (PLL-MPs). Both the number of macrophages with particles as well as number of particles being internalised by Mtb-infected cells drastically increased within 2 h of particle incubation. Previous studies have also shown that particles with increasing positive surface charge show corresponding increase in uptake by mouse macrophages and primary human antigen presenting cells [26, 34, 35]. This is observed because mammalian cells have sialic acid, anionic proteoglycans, and glycosaminoglycans embedded throughout the plasma membrane that can interact electrostatically with positively charged particles [36, 37].

To investigate further, we encapsulated rif in MPs and notable increase in intracellular rif concentration was observed. rif loaded PLL-MPs performed significantly better than nonmodified particles in eliminating intracellular H37Rv. Clemens *et al.* explored rif loaded Mesoporous Silica Nanoparticles coated with cationic polymer Polyethyleneimine [38]. These 100 nm sized nanoparticles with positive surface charge demonstrated enhanced killing of intracellular Mtb compared to non-coated and free drug. However, the size range of these particles (~100 nm) was not in the inhalable size range for deep lung deposition through inhalation [39–41].

In this regard, we have explored the size range that is known to deposit in deep lungs after dry powder delivery (500 nm to 2 μm). Ohashi *et al.* showed that 1 μm PLGA particles deposit in the lungs initially but begin to localize in the trachea over time, due to mucociliary clearance [42]. Hence, it is important to ensure that the particles get rapidly internalized in the target cells to avoid clearance. Immune cells, particularly AMs, are major host cells for Mtb and inhalation ensures that the drug loaded MPs get phagocytosed by the infected cell to achieve high drug concentrations in the bacterial niche [43–45]. While inhalation-based delivery inherently targets AMs, we have shown that prompt uptake of cationic particles by AMs can result in two-fold higher uptake in mice, thus enhancing delivery efficacy and preventing loss of drug. To our knowledge, this is the first report demonstrating improved delivery to AMs *in vivo* using microparticles with cationic surface coating.

Inhalable particle formulations also improve drug pharmacokinetics. Sustained release MPs deposit in the deep lung, where they form a depot, allowing initial high drug concentration in the lung, which then diffuses systemically [14]. Free rif delivered via inhalation to guinea pigs was undetectable in serum after 3 h, while drug released from its polymeric formulation was detected till 8 h [46]. Delivery of MPs through inhalation route enables upto 4 times longer systemic half-life compared to free drug administered via intravenous route, hence therapeutic drug concentrations are maintained for extended duration, as shown in healthy mice [16]. In rhesus macaques, inhalable MPs containing anti-TB drugs were delivered through dry powder inhalation and resulted in higher concentration of drug in lung compared to liver and kidneys, along with 1.5-4 times longer serum half-life of antibiotics juxtaposed to intravenous delivery of free drug [17].

Although positively charged particles can cause toxicity by destabilizing plasma membranes [47], both polymers PLGA and poly-L-lysine used in this study are US FDA approved. PLGA is used extensively in injectable depot formulations (Atridox® for doxycycline, Ozurdex® for dexamethasone etc. [48]) and Poly Lysine has been given GRAS status and is used as a food preservative [49]. We had also confirmed that microparticles formed using these polymers were cytocompatible with THP-1 macrophages. Additionally, PLL-MPs exhibited negligible local lung toxicity as inferred from histological studies in mice.

Furthermore, microparticles can be used as targeted delivery vehicles by incorporating specific ligands or by using smart polymers that can target specific cells, sense microenvironmental cues and cause triggered release [38, 50–53]. Hydrophilic anti-TB drugs like aminoglycosides and INH have been encapsulated in micro-carriers and have displayed improved intracellular delivery [54, 55]. Albumin microcarriers have been used to improve solubility and bioavailability of antimycobacterial benzothiazinone compounds [56]. Ligands such as mannose and fucose were used for targeted intracellular delivery into AMs that express high levels of mannose receptors [57–60]. Such strategies can be further leveraged to improve the delivery efficiency of positively charged particulate systems.

The effect of cationic particle delivery remains to be tested on an *in vivo* model of TB infection. This will require testing of cationic particles in animal BSL-3 facilities over 2-3 months after infection and will be done in future studies. Delivery of microparticles through the granuloma is challenging due to its size and low diffusion, however, sustained release formulations could still allow high local concentration of released drugs that can permeate through the granuloma. Some studies have shown accumulation of liposomes and polymersomes within the granuloma in *M. marinum* infected zebrafish and in *M. tuberculosis* infected mice after intravenous administration [61, 62]. Thus, microparticle-based targeted delivery to Mtb-infected macrophages has the potential to improve existing TB therapy and should be further explored.

## 5. Conclusion

In this study, we examined how physical parameters of microparticles can be modulated to enhance delivery into Mtb-infected cells.

1. We observed that Mtb-infected cells were highly phagocytic and PLGA particles of all three sizes 500 nm, 1 μm and 2 μm were internalized favourably by infected macrophages.
2. Modification of surface charge by conjugating cationic polymer Poly-L-Lysine resulted in significantly more Mtb-infected cells internalizing these particles in large numbers *in vitro* which also translated in higher bactericidal efficacy compared to free drug and non-modified particles.
3. The cationic particles also improved delivery to immune cells and specifically alveolar macrophages *in vivo* after pulmonary delivery.
4. Our study shows that particle engineering approaches enable improved drug delivery to achieve desired effects and inhalation-based delivery of cationic particles can be effective for the treatment of TB.

## 6. Acknowledgements

We acknowledge intramural support from the Indian Institute of Science to Rachit Agarwal. Extramural support for funding this research was received from Department of Science and Technology (DST, India) Ramanujan Fellowship (SB/S2/RJN-036/2017), and Mr. Lakshmi Narayanan. This work was supported, in part, by the Bill & Melinda Gates Foundation [OPP1210498]. Under the grant conditions of the Foundation, a Creative Commons Attribution 4.0 Generic License has already been assigned to the Author Accepted Manuscript version that might arise from this submission. We also acknowledge stipend support to Pallavi Raj Sharma from Department of Biotechnology (DBT, India) Junior Research Fellowship program (DBT/2017/IISc/941). Funding from Dr Vijaya and Rajagopal Rao for Biomedical Engineering research at the Centre for BioSystems Science and Engineering is also acknowledged. We thank Prof. Deepak Saini, IISc (Bengaluru, India) and Prof. Lalita Ramakrishnan, University of Cambridge (Cambridge, UK), for providing the GFP expressing mycobacterial strain H37Rv. Prof. Amit Singh, IISc (Bengaluru, India) is duly acknowledged for providing THP-1 cell line and H37Rv wild type strains. Central animal facility, BSL-3 facility, AFMM facility IISc, Biological Sciences Imaging Facility, BSSE Central Facility, Materials Engineering Central Facility and MRDG Central Facility are acknowledged for access to various instruments. We are thankful to Dr Siddharth Jhunjhunwala (IISc, Bengaluru) for his assistance with flow cytometry panel design and data analysis.

