## Supplementary Information for "Cationic inhalable particles for enhanced drug delivery to *M. tuberculosis* infected macrophages"

**Supplementary Methods**

### 1. Surface charge modification using EDC-NHS chemistry

To modify the surface charge, 2 mg mL$^{-1}$ PLGA particles were suspended and mixed on a rotor with 4 mM EDC and 5 mM NHS in 0.1 M MES buffer (pH 5.6) for 2 h followed by 4 h incubation with 0.1% poly-L-lysine in 0.1 M MES buffer (pH 7.4). The particles were then washed thrice with deionized water, snap-frozen and lyophilized.

### 2. Size and charge determination

Microparticle size and charge were characterized using Malvern Nano ZS ZetaSizer (Malvern Instruments, Worcestershire, UK). The lyophilized microparticle sample was weighed (1 mg) and suspended in deionized water to obtain a homogenous solution. The sample was sonicated in a bath sonicator to ensure that there are no clumps before acquiring data using dynamic light scattering (DLS). Zeta potential measurement was performed using folded capillary zeta cell (Malvern DTS 1070).

### 3. Scanning Electron Microscopy

To observe particle morphology, scanning electron microscopy (SEM) of the samples was carried out. Particles of all sizes, with poly-L-lysine coating and with encapsulated rifampicin were suspended in distilled water (0.5 mg mL$^{-1}$) and briefly sonicated. 10 μL of sample was distributed evenly on a double-sided carbon tape placed on a metal stub and vacuum dried, followed by sputter coating with gold (20 nm) and rastered using an electron beam (5 keV).

### 4. WST-8 Assay

Cytocompatibility of the particles was assessed using WST-8 assay. Briefly, PMA treated THP-1 cells were seeded in 96 well plates and incubated with 50 μg mL$^{-1}$ and 500 μg mL$^{-1}$ particles resuspended in 1x PBS. After 24 h, WST-8 developer reagent and electron mediator solution were mixed and added and allowed to incubate for 2 h. Absorbance readings were taken at 450 nm.

5. **Murine Bone Marrow Derived Macrophages (BMDMs) isolation and culture**

C57BL/6 WT mice (8 – 10 weeks) were euthanized and the femur and tibia were dissected and treated with 70% ethanol for 5 min. In a sterile environment, the ends of the bones were cut and media was passed through the hollow bone using a 26 G needle into a sterile petridish. The collected cells were passed through a 19 G needle to break clumps. After RBC lysis, cells were counted and seeded into 12 well plates at a seeding density of 2 x $10^6$ cells/well with 20 ng $mL^{-1}$ M-CSF. Every alternate day, half the volume of media (500 µL) was removed and replenished with fresh media containing M-CSF (20 ng $mL^{-1}$). On day 7, cells were washed and incubated in fresh media. BMDMs were characterized by assessing expression of CD11b and F4/80 surface markers using flow cytometry.

6. **Flow cytometry**

Particles suspended in 1X PBS (2 mg $mL^{-1}$) were briefly sonicated and added to uninfected or infected PMA treated THP-1 or BMDMs at a final concentration of 50 µg $mL^{-1}$. After 4 h (or otherwise mentioned) incubation, cells were washed thrice with 1x PBS and scraped out of the wells. For H37Rv infected cells, Fixable Violet Live Dead Dye was used to stain dead cells and cell suspension was fixed using 4% Paraformaldehyde. For antibody staining, cells were incubated with antibody and respective isotype for 30 min at 4 °C. Finally, cells were suspended in FACS Buffer (1% BSA in 1X PBS-EDTA) and data was acquired using flow cytometry (BD FACS Celesta). Data was analysed using FlowJo v9. **Supplementary figure 3** shows the gating strategy to identify live cells, Mtb-infected cells and cells with particles.

7. **Confocal imaging**

THP-1 monocytes were seeded on gelatin (100 µg $mL^{-1}$) coated cover slips followed by PMA differentiation and infection with mycobacterial strain GFP-H37Rv. After incubation with microparticles, the cells were washed thrice with 1x PBS and fixed using 4% Paraformaldehyde. ActinGreen$^{TM}$488 ReadyProbes$^{TM}$ reagent was used to stain the cytoskeleton and nuclear staining was done using Hoechst 33342 stain (1 µg $mL^{-1}$). The coverslips were mounted on slides using glycerol, dried and sealed. The images were acquired using Zeiss LSM880 (Airyscan) at 63x magnification with oil and analysed using ImageJ.

8. **Quantification of intracellular rifampicin using HPLC:**

Rif was quantified using high-pressure liquid chromatography (HPLC, Prominence-i LC-2030 Plus, Shimadzu). Briefly, standard solutions of rif were injected in a C18 column attached to the HPLC and eluted under a gradient flow of methanol and water at 0.1 mL $min^{-1}$. The absorbance was measured at 280 nm. The corresponding peak for rif on the chromatogram was identified, and its AUC was calculated using Shimadzu LabSolutions software. This AUC was used to generate a standard curve of rif. Intracellular rif concentration was determined by extracting intracellular components of treated cells and quantifying using HPLC. Rif and rif-loaded microparticles were incubated with PMA differentiated THP-1 macrophages (2 x $10^5$ cells per well) for 4 h. The cells were washed twice with PBS, scraped, and collected in 1.5 mL tubes. The cell suspension was frozen and thawed thrice to cause cell lysis. The lysates were lyophilized and dissolved in 100 µL of 50% methanol and quantified using HPLC.

9. **Drug release analysis**

For drug release experiment, 10 mg $mL^{-1}$ of rif-MPs were incubated in 1x PBS (pH 7.4) on a rotor maintained at 37 °C. At each time point, the tubes were centrifuged at 8000 g (10 min) and the pellet was resuspended in 100 µL DMSO. The concentration of rif remaining in the particles was determined by measuring absorbance at 335 nm.

**10. Cryosectioning**

For cryosectioning, mice were euthanized after intratracheal administration of particles. 1 mL of Polyfreeze and 4% paraformaldehyde (PFA) solution (mixed in 1:1 ratio) was flushed through the trachea to inflate the lungs. After 30 min, the trachea was sealed with a thread, lungs were extracted and embedded in Polyfreeze solution. The sample was frozen at -20 °C and 5 µm sections were cut using a cryotome (Leica CM 1510 S). The sections were dried and Polyfreeze was dissolved by incubating in 1x PBS for 1 h. The sections were stained with Hoechst 33342 stain (1 µg $mL^{-1}$) and imaged under an upright fluorescence microscope (Olympus BX53F).

**11. Histological analysis**

Post-intratracheal delivery of PLGA particles, mice were sacrificed after 1 and 7 d. Lungs were inflated with 2% PFA and stored overnight. Following wash with MilliQ water, lungs were processed and embedded in paraffin wax. Blocks were trimmed and kept on ice overnight followed by sectioning (5 µm sections) using Leica HistoCore MULTICUT and collected on poly-L-lysine coated glass slides. The sections were rehydrated with a series of ethanol gradients followed by staining with Mayer’s hematoxylin (4 min) and Eosin Y (45 sec) with intermittent washing.

### Supplementary Figures

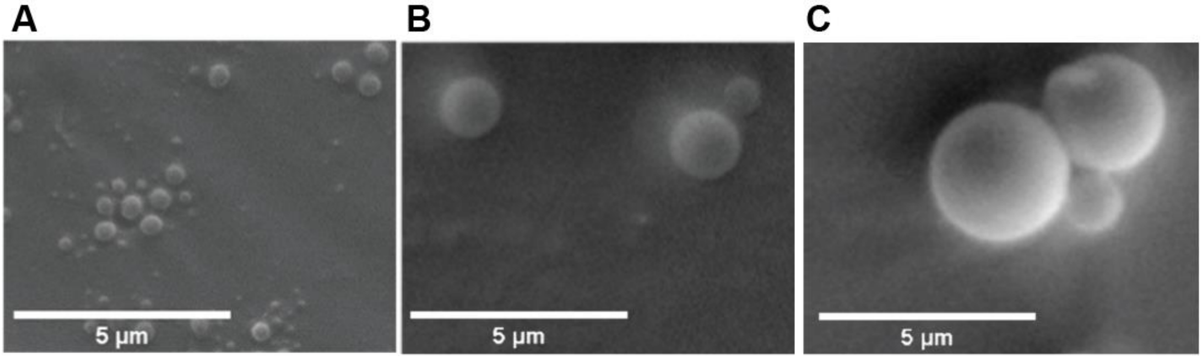


**Supplementary figure 1. PLGA MPs had smooth and round morphology.** Scanning Electron Microscopy (SEM) images of PLGA particles of (A) 500 nm, (B) 1 µm, and (C) 2 µm diameter.

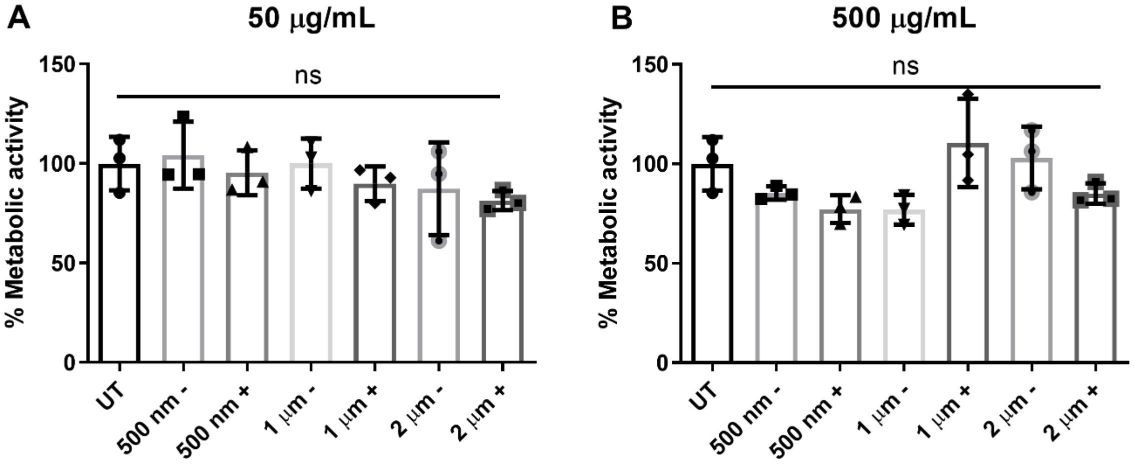


**Supplementary figure 2. PLGA microparticles of different sizes and charges are cytocompatible.** Percentage of metabolic activity of THP-1 macrophages reported after 24 h incubation of particles at (A) 50 µg $mL^{-1}$ and (B) 500 µg $mL^{-1}$ concentration. Values were normalized to the values corresponding to the wells in which no particles were added (Untreated). All groups were found statistically non-significant (ns) using ordinary one-way ANOVA followed by Tukey's multiple comparisons test. Data were represented as mean ± SD.

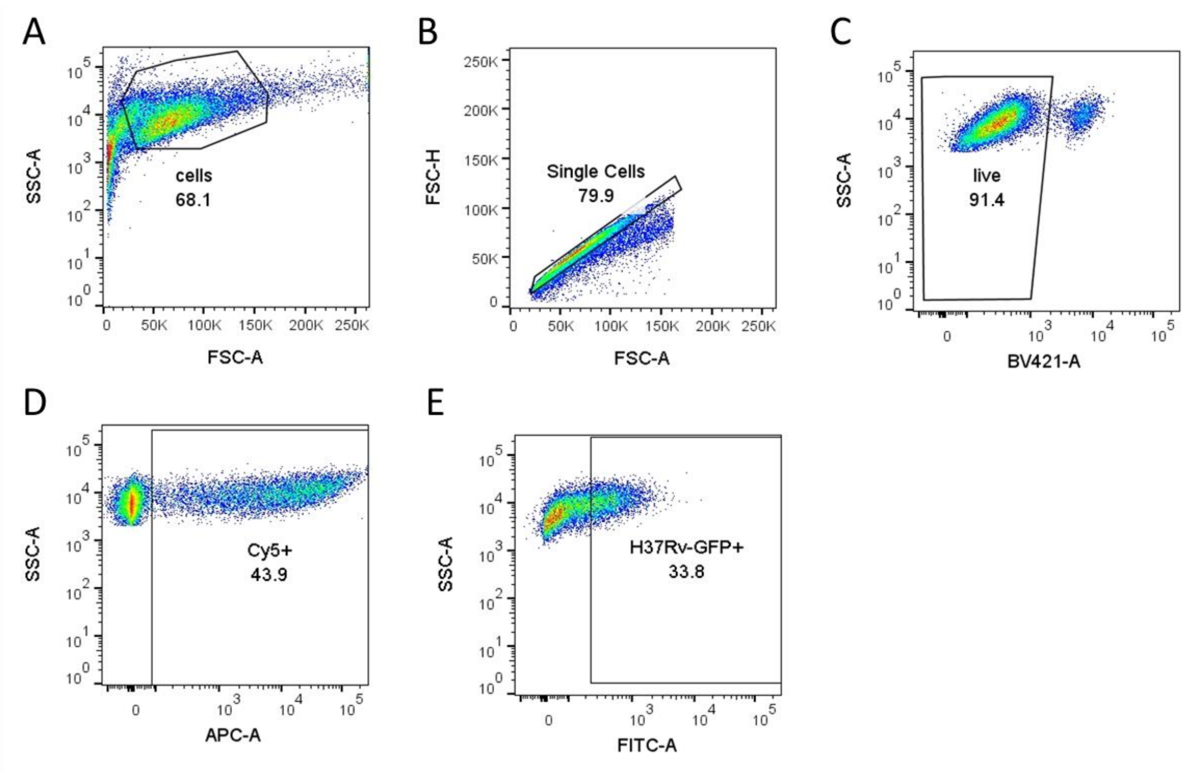


**Supplementary figure 3**. **Representative gating strategy to identify H37Rv-GFP infected cells and cells with microparticles**. (A) Forward scatter (FSC-A) vs Side scatter (SSC-A) was used to identify cells. (B) Within the cells gate, single cells were identified by plotting FSC-H vs FSC-A. (C) Fixable Live-dead violet staining was done to identify live cells. (D) Cells in APC channel used to identify Cy5 positive population, i.e., cells with particles. (E) Infected cells in FITC channel gated to identify GFP positive population, i.e., cells with bacteria.

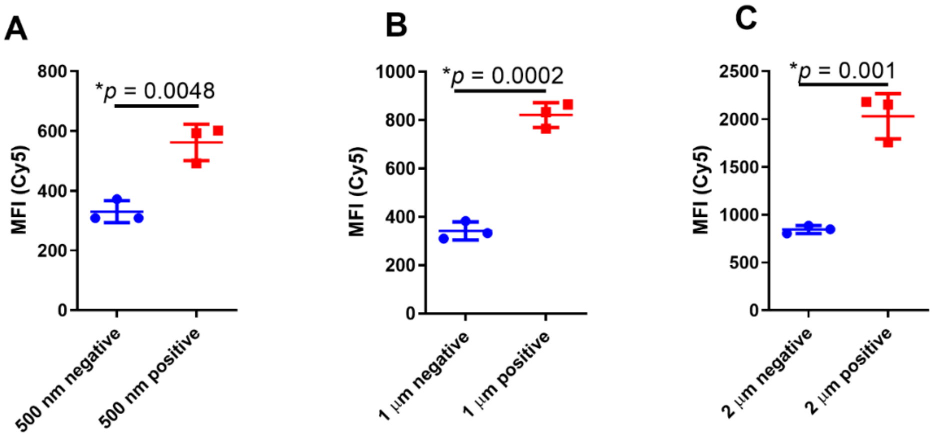


**Supplementary figure 4**. **Cationic microparticles accumulate in higher numbers in THP-1 macrophages.** Median Fluorescence Intensity (MFI) values reported for Cy5 channel (negatively charged particles in blue circles and positively charged particles in red squares) in PMA- differentiated THP-1 macrophages incubated with (A) 500 nm, (B) 1 µm and (C) 2 µm of both negative and positive surface charge. Two-tailed t-test was used to determine significance. All groups have n = 3 and data were represented as mean ± SD.

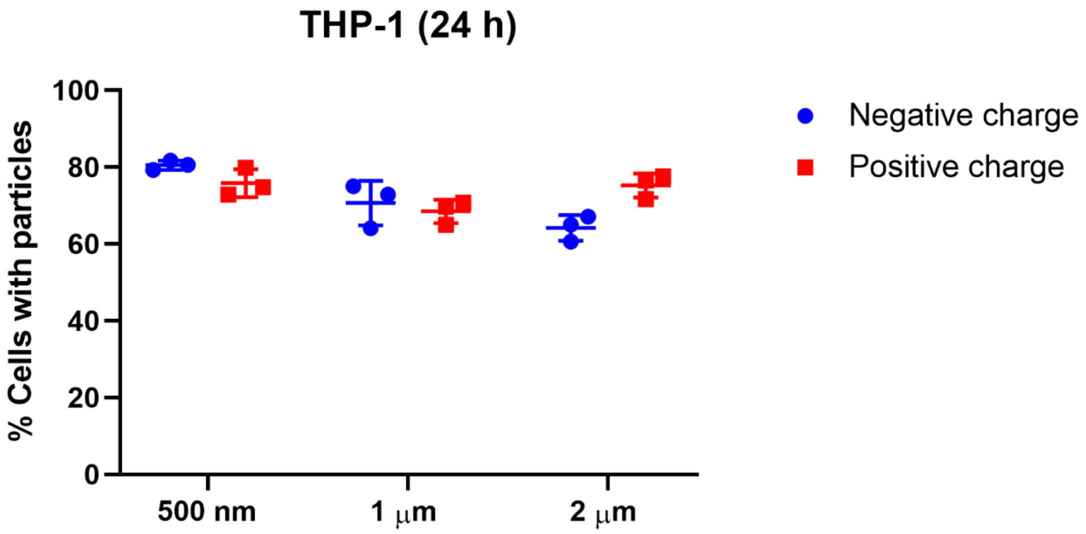


**Supplementary figure 5. THP-1 macrophages uptake of MPs is saturated by 24 h.** Percentage of THP-1 macrophages with PLGA particles of different sizes and charges reported after 24 h incubation. Data were represented as mean ± SD.

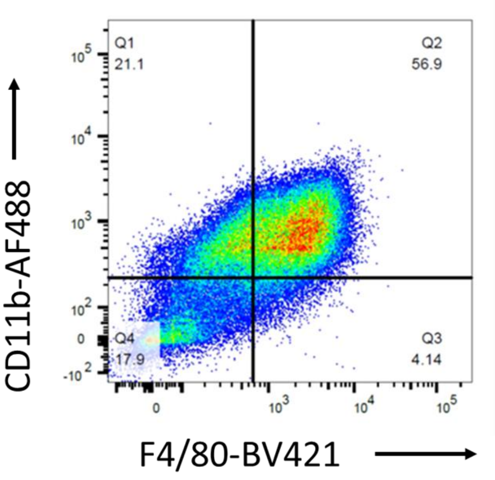


**Supplementary figure 6. Gating strategy and purity of bone marrow derived macrophages (BMDMs).** Murine primary BMDMs were analysed for surface marker expression of CD11b and F4/80 to determine purity.

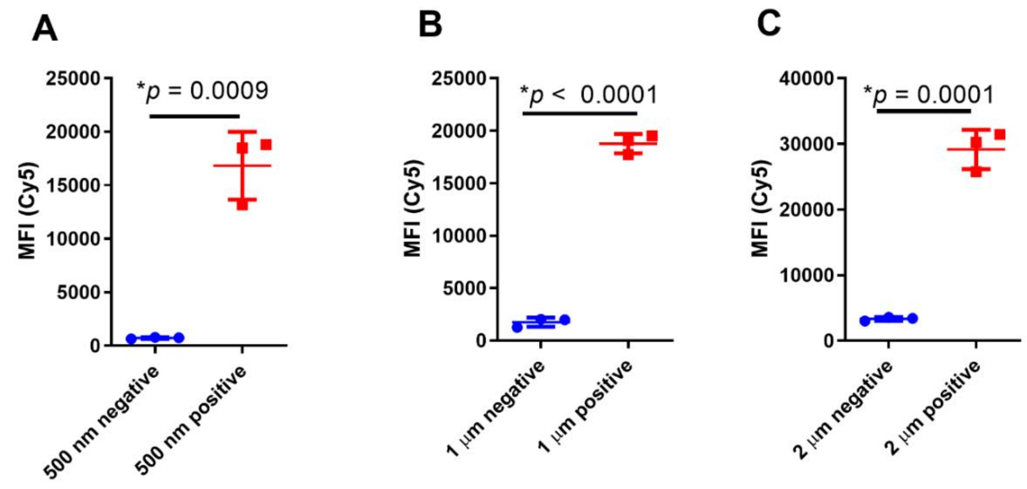


**Supplementary figure 7. Cationic microparticles accumulate in higher numbers in BMDMs.** Median fluorescence intensity (MFI) (negatively charged particles in blue circles and positively charged particles in red squares) plotted for murine Bone Marrow Derived Macrophages incubated with (A) 500 nm particles, (B) 1 µm particles and (C) 2 µm particles. Analysis was done using two-tailed t-test and all groups had n = 3. Data were represented as mean ± SD.

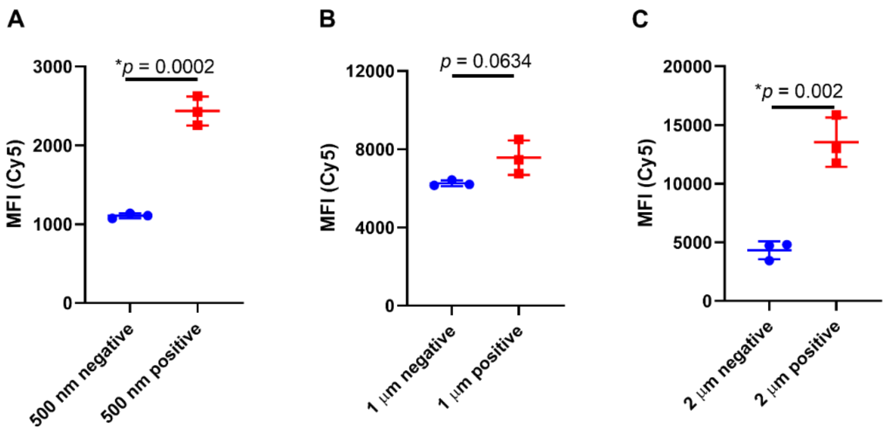


**Supplementary figure 8. Cationic microparticles accumulate in higher numbers in H37Rv infected THP-1 macrophages.** Representative plots of median fluorescence intensity (MFI) (negatively charged particles in blue circles and positively charged particles in red squares) for H37Rv infected THP-1 macrophages incubated with (A) 500 nm particles, (B) 1 µm particles and (C) 2 µm particles. Analysis was done using two-tailed t-test and all groups had n = 3. Data were represented as mean ± SD.

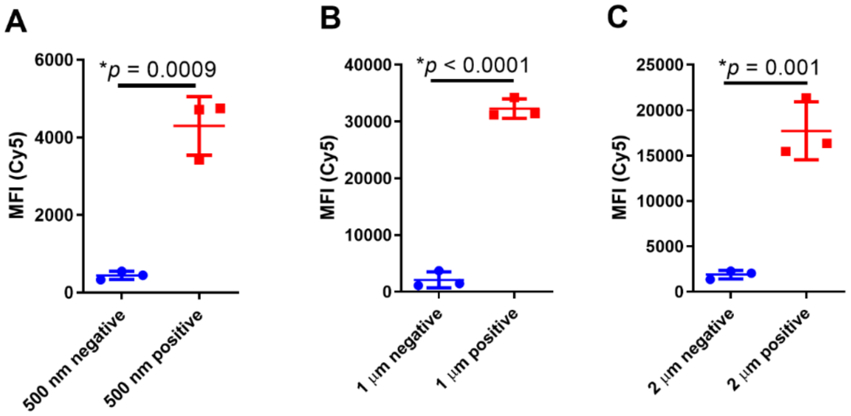


**Supplementary figure 9. Cationic microparticles accumulate in higher numbers in H37Rv infected BMDMs.** MFI plotted (negatively charged particles in blue circles and positively charged particles in red squares) for H37Rv infected BMDMs incubated with (A) 500 nm particles, (B) 1 µm particles and (C) 2 µm particles. Analysis was done using two-tailed t-test. All graphs had n = 3. Data were represented as mean ± SD.

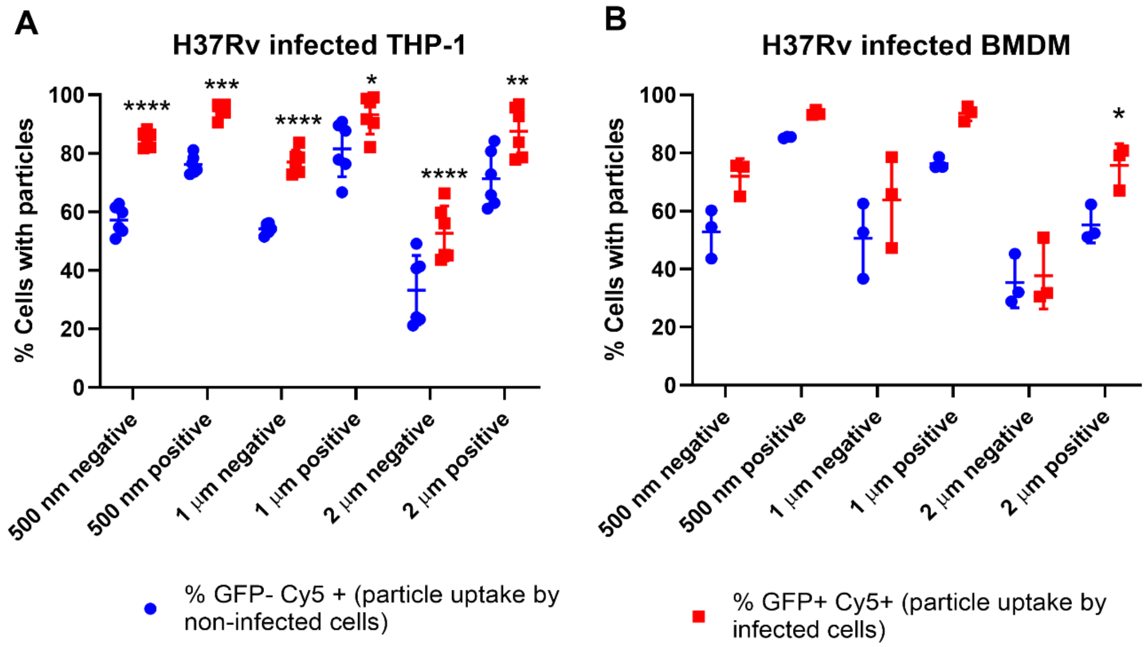


**Supplementary Figure 10. H37Rv-infected macrophages exhibit enhanced particle uptake.** Percentage of cells with particles is plotted for H37Rv infected (GFP positive, red squares) and non-infected cells (GFP negative, blue circles) in (A) THP-1 macrophages (n = 6, **p* = 0.0314, ***p* = 0.001, ****p* = 0.0002, *****p* < 0.0001) and (B) BMDMs (n = 3, **p* = 0.0368). Data were analysed using two-way ANOVA followed by Sidak's multiple comparisons test. Data were represented as mean ± SD.

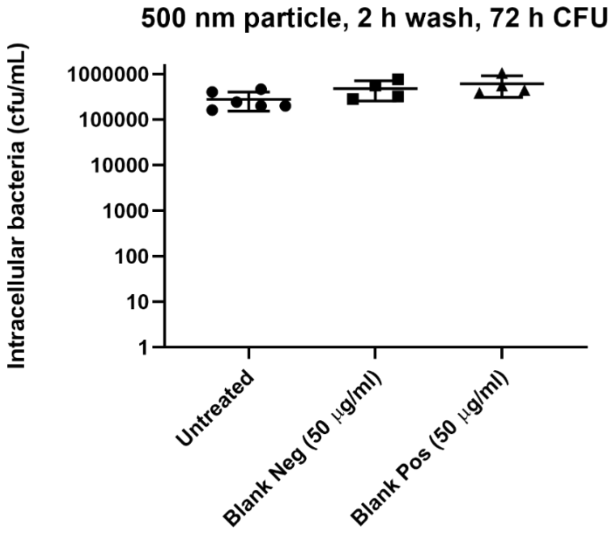


**Supplementary figure 11. Blank microparticles had no effect on intracellular CFU.** Intracellular bacterial counts (cfu/mL) from H37Rv-infected THP-1 macrophages reported after 2 h treatment with 500 nm microparticles – non-modified (Neg) and poly-l-lysine conjugated (Pos). Ordinary one-way ANOVA followed by Tukey's multiple comparison test was used to determine statistical differences. All groups were found non-significant. Data were represented as mean ± SD.

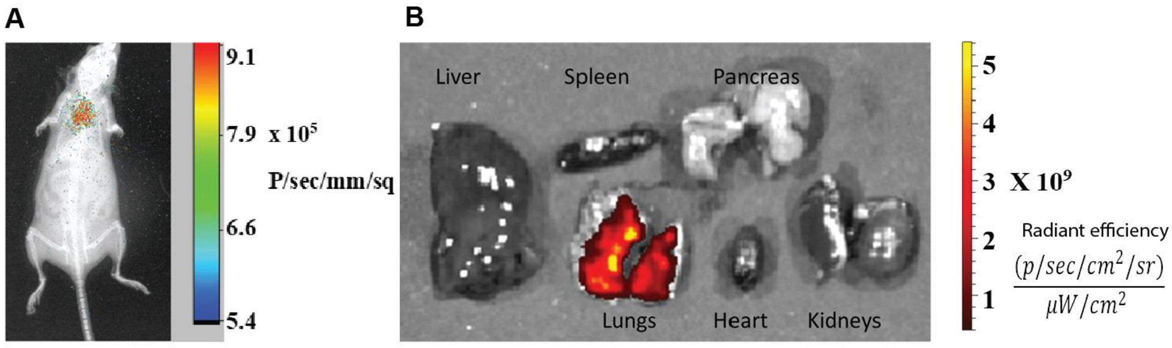


**Supplementary figure 12. PLGA microparticles deposited in lungs after intratracheal instillation.** (A) In vivo imaging of healthy mouse using In Vivo Xtreme II (Bruker®) after intratracheal delivery of microparticles. X-ray image was overlayed with fluorescence and scale bar denotes radiance efficiency. (B) Ex-vivo fluorescence imaging of explanted organs using In Vivo Xtreme II (Bruker®) after intratracheal MP delivery.

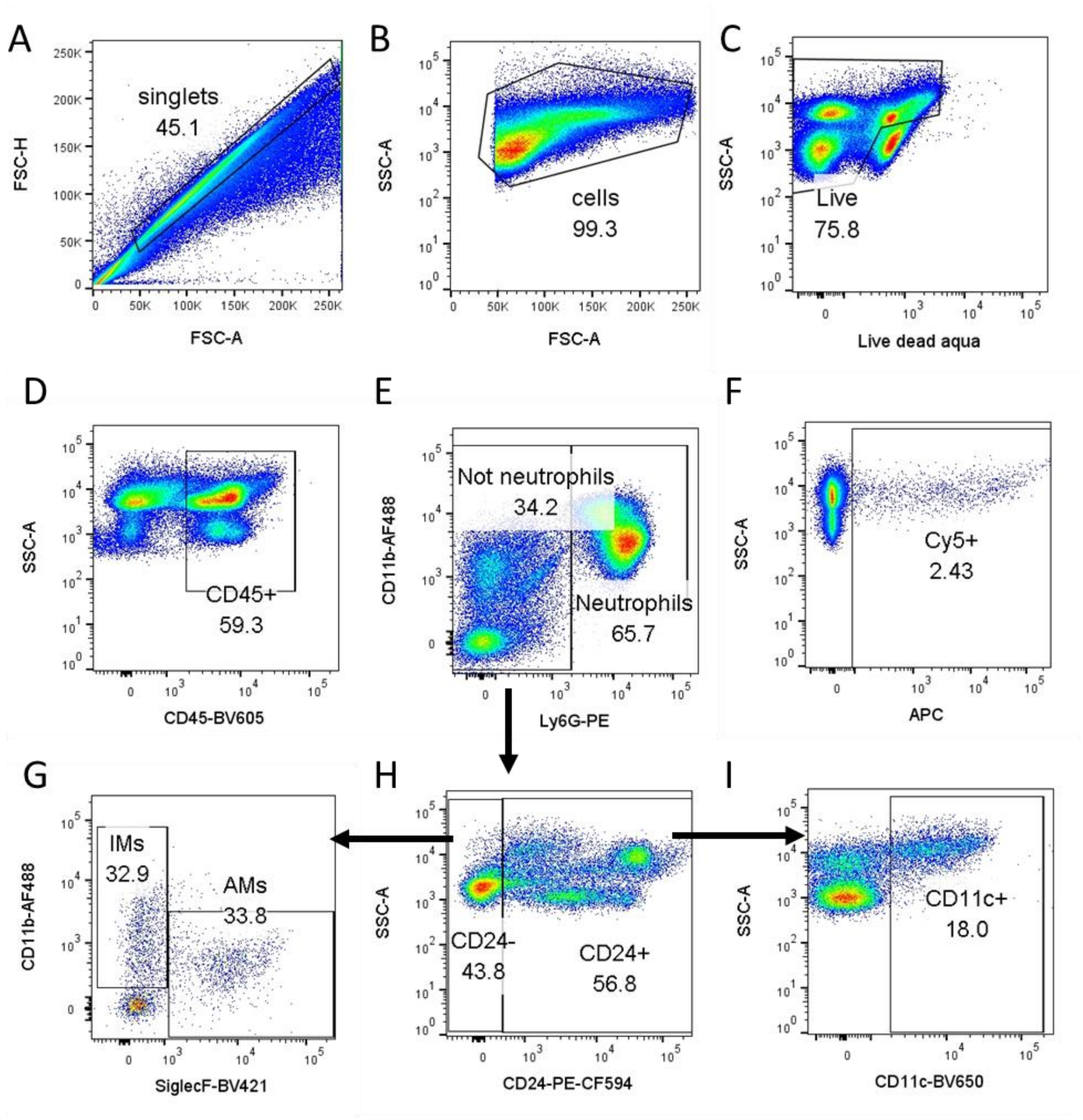


**Supplementary figure 13. Gating strategy to identify different immune cell populations in the lung.** Mice were sacrificed 1.5 h after intratracheal particle delivery. (A) Debris was removed by singlet gating and (B) cells were gated from FSC vs SSC plot. (C) Live cells were gated using FMO (Fluorophore Minus One). (D) Primary fluorophore conjugated antibodies were used to identify immune cells (CD45+), all further gating was done on CD45+ subset. (E) Neutrophils were gated using CD11b+ Ly6G+, and “Not Neutrophil” (Ly6G-) subset was used for further gating. (H) CD24 was used to distinguish between Dendritic cells (DCs) and macrophages. (G) CD24- subset was then used to identify alveolar macrophages (CD24- CD11b- SiglecF+) and interstitial macrophages (CD24- CD11b+ SiglecF-). (I) Dendritic cells were identified by CD11c+ gating in CD24+ subset (Ly6G- CD24+ CD11c+). (F) Particles were detected in the APC channel. Compensation beads were used as single colour controls for all antibodies and respective FMOs were also prepared and used for gating.

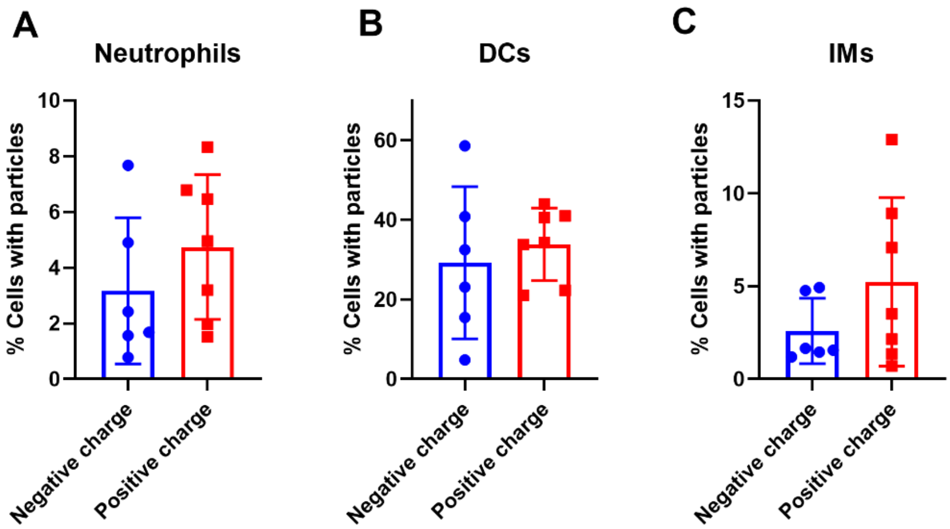


**Supplementary figure 14. Cationic particles show increased trend in particle uptake in different immune cell populations after pulmonary delivery.** Non-modified-MPs (negative charge) and PLL-MPs (positive charge) were delivered to mice lungs and sacrificed after 1.5 h. Flow cytometric plots showing particle uptake in (A) neutrophils, (B) dendritic cells (DCs), and (C) interstitial macrophages (IMs) was plotted. Two tailed t-test was used to determine significance and differences between the groups in all cell types were found non-significant. Data were represented as mean ± SD.
